# Bottlenose dolphins’ broadband clicks are structured for communication

**DOI:** 10.1101/2023.01.11.523588

**Authors:** Arthur Stepanov, Hristo Zhivomirov, Ivaylo Nedelchev, Penka Stateva

## Abstract

Bottlenose dolphins’ broadband click vocalizations are well studied in the literature with respect to their echolocation function. Their use for communication among conspecifics has long been speculated, but not conclusively established so far. In this study we categorize dolphins’ click productions into types on the basis of their amplitude contour and analyze the distribution of individual clicks and click sequences against their duration and length. We demonstrate that the repertoire and composition of clicks and click sequences follow three key linguistic laws of efficient communication, namely, Zipf’s rank-frequency law, the law of brevity and Menzerath-Altmann law. Conforming to the rank-frequency law suggests that clicks may form a linguistic code that is subject to selective pressures for unification, on the one hand, and diversification, on the other. Conforming to the other two laws also implies that dolphins use clicks in accord with the compression criterion, or minimization of code length without loss of information. Our results furnish novel evidence for conformity to the linguistic laws in this type of dolphins’ signal and in the realm of animal vocalizations more generally.

## Introduction

Two of the most studied types of acoustic signals produced by bottlenose dolphins (*tursiops truncatus*) are whistles and clicks. Whistles are typically produced in the frequency range that is accessible to human hearing and are often discussed in the context of communication [32]. Clicks are short broadband pulses of vibration produced by a pair of nasal lips located right beneath the blowhole in the upper part of the dolphin’s head; the emitted sound then propagates through the lipid mass of the dolphin’s melon into the narrow-directional signal beam outwards into the water [1][2][3]. Clicks are typically produced as separate pulses but also in trains or sequences in a broadband frequency range between 20÷150kHz (op.cit.), far beyond the frequency range for hearing and sound producing capacity of humans (see Figure 1).

**Figure 1.**
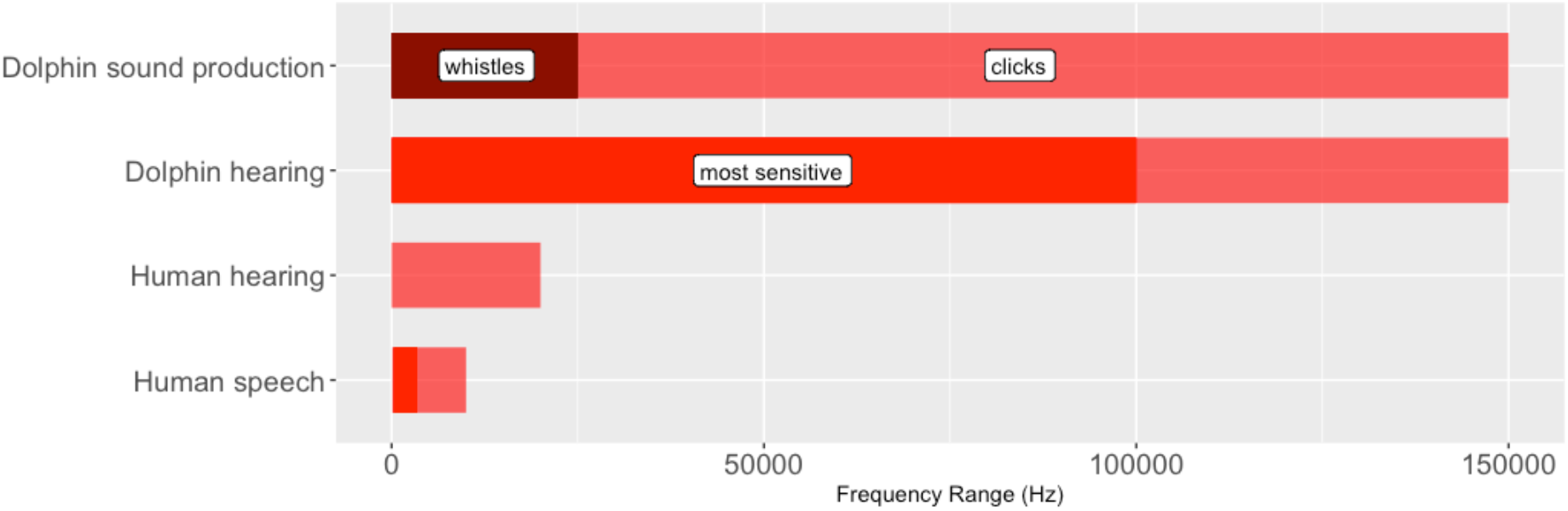
Comparison of frequency ranges pertaining to human and dolphin hearing and sound production (e.g. [35])

Dolphins’ use of clicks for echolocation is widely documented in the literature. Similarly to whistles, click sequences have been increasingly often implicated for communication among conspecifics at various distances [4][5], although no controlled study has directly supported this conjecture so far.

The communicative aspect of signaling in various animal species is commonly evaluated through a degree of conforming to the so called »linguistic laws« of efficient communication, a series of observed regularities in the communicative use of linguistic code that go back to Zipf’s influential work [6] and have been further elaborated in the context of information theory [7]. The distribution of whistles, another well studied type of acoustic signal produced by bottlenose dolphins, has been shown in a controlled study to conform well to predictions of the Zipf’s frequency-rank law (see below). In this report, we explore the distribution of click and click sequences produced by adult bottlenose dolphins living in captivity and demonstrate that it properly conforms to three linguistic laws: i) Zipf’s frequency-rank law, ii) the law of brevity and iii) Menzerath-Altmann law that relate frequency of usage and frequency rank and explore the connection between sequence length and duration of specific sequence components.

## Linguistic laws

### Zipf-Mandelbrot’s rank-frequency law

Perhaps the most widely known of the family of linguistic laws popularly known as Zipf’s law but is only one of many laws in Zipf’s popular book [6].

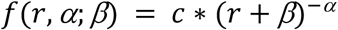

where *r* is a frequency rank of a word, and *f* is its frequency in a natural corpus. α is a positive parameter, *β* is Mandelbrot’s “rank-shifting” constant (which was not there in Zipf’s original formulation of the law). *α ≈* 1 and *β* = 0 for English words. The law formalizes a negative correlation between *f* and *r* is for α > 0 which converges to a linear dependency with a negative slope on a logarithmical scale. In other words, the most frequent English word (*r=1*) has a frequency proportional to 1, the second, third etc. most frequent words (*r*=2,3,…n) has a frequency proportional to 1/2, 1/3, … 1/n of the frequency of the first most frequent word. For word systems for which *α* ≠1 the proportional coefficient will be 1/2*α*, 1/3*α* and so on. Zipf’s law has been observed in many different levels of structural organization of natural language, but also in a wider contexts of biological organization of living systems, in particular at the intra-molecular level (codons and DNA coding) as well as at the macro levels such as larger ecological systems (see [34] for areview of different forms of Zipfian dependencies and a comparative analysis of their applicability across various linguistic and biological domains).

This reverse correlation is often thought to arise as a result of selectional tension between two opposing functional tendencies: that for unification of the linguistic code (resulting, in the limit, in a single code for conveying all possible messages, or ultimate repetition, which places most burden on the decoder, or the hearer, to deduce information), on the one hand, and its diversification (aiming, ultimately, for a unique and non-repeating code for each message, thus no repetition at all, which places most burden on the speaker), on the other. In the realm of human communication, both speakers and hearers strive to minimize their respective effort thus leading to some sort of equilibrium between the two functional tendencies in the communicative system. In a so optimized communication, then, an inverse correlation between frequency and rank results in a linear dependency slope of around −1.00 for human language, on a log-log scale [7].

In the realm of dolphin cognition research, [8] [9] [10] used a variety of k-means clustering technique to categorize whistles produced by adult and infant bottlenose dolphins into discrete types and analyzed their distribution with respect to Zipf’s rank-frequency law. According to these studies, this distribution manifests a linear dependency on a log-log scale with a slope varying from about −0.82 for newborns then reaching up to −1.07 within the period of 2÷8 months and converging on the adult value of about −0.95 afterwards. This indicates that dolphins’ use of whistles is broadly in line with the predictions of the least effort principle and the functional tension hypothesis. It is therefore reasonable to suppose that similar considerations may also apply regarding dolphin click signals.

### Zipf’s “law of abbreviation”

Zipf’s “law of abbreviation” broadly states that the length of a linguistic unit reversely correlates with its frequency of use, viz. more frequent sequences tend to be shorter. Zipf himself did not specify a formal relationship between frequency and length. [11] looked in more detail at the relationship between word length in terms of letters in English and Swedish and their frequencies, and also the length of sentences in terms of number of words in the Brown corpus of written American English texts [12]. They found that the relationship is best modeled via a version of a gamma distribution whose probability density is described below:

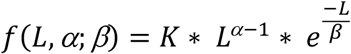

where L is the sequence length, *α* the ‘shape’ parameter determining the existence and the location of the function peak, and *β* the ‘scale’ parameter determing the ‘spread’ of the distribution. Because this law is somewhat less known that the rank-frequency law, we illustrate it here with a few examples. Figure 2 illustrates the distributions of the predicted and actually observed frequencies in the Brown corpus of written American English texts.

**Figure 2.**
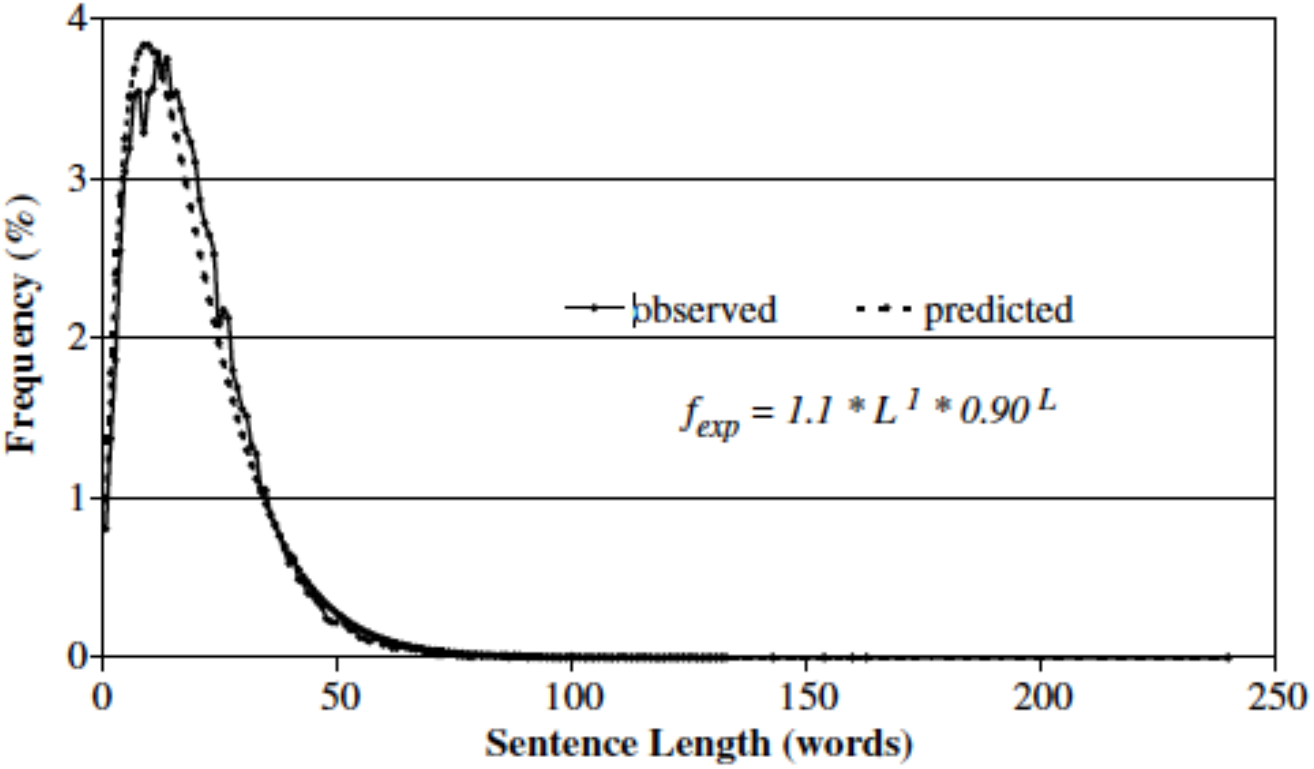
Observed and predicted sentence frequencies on the basis of sentence length in words (L) in the entire Brown corpus of written American English texts. Reproduced from Sigurd et al. (2004) with the publisher’s permission.

Perhaps even more relevant to our purposes, a similar distribution has been shown to obtain in the realm of spoken speech. [13] conducted a quantitative analysis of sentence lengths in the inaugural speeches of US presidents between 1789 and 2021 and the annual speeches of UK party leaders between 1895 and 2018. They argue that the distribution of sentence length over frequencies obeys Weibull distribution as a consequence of the Least Effort principle attempting to minimize the effort for expression of thoughts. Two examples of the observed frequency distributions are illustrated in Figure 3.

**Figure 3.**
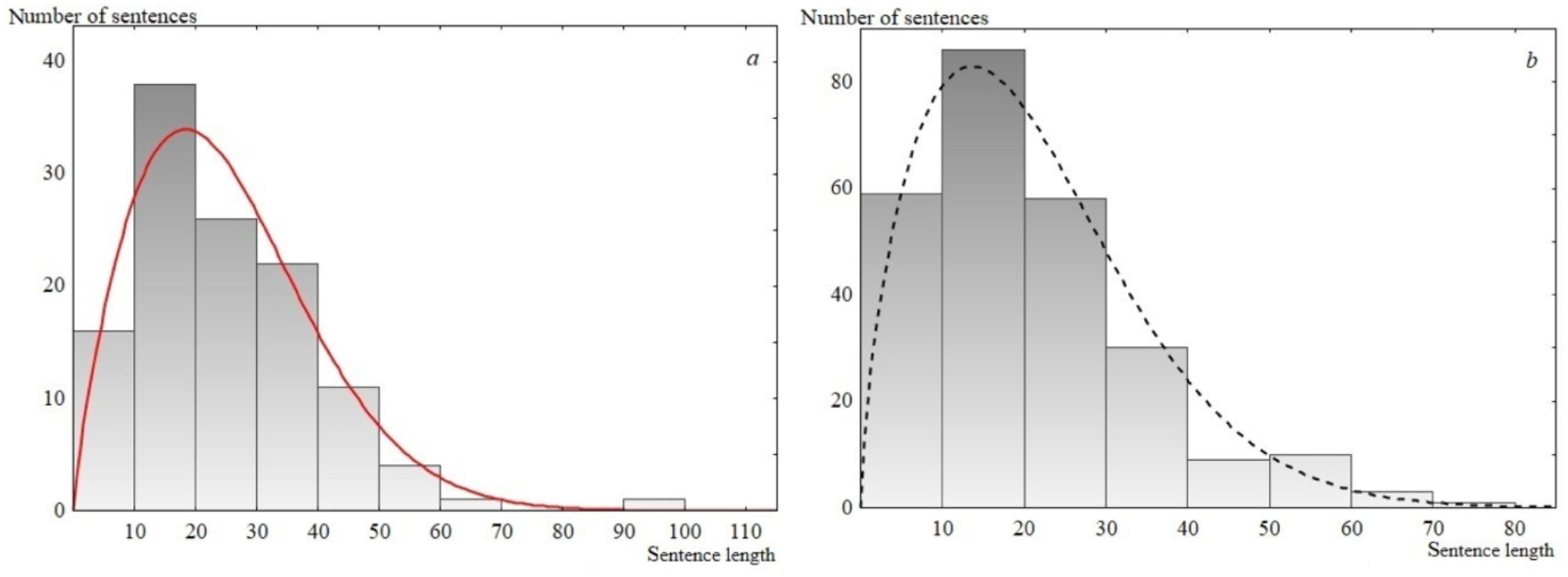
Sentence length distribution histogram. (a) USA inaugural presidential speech, 1817. The solid line is the Weibull distribution. (b) UK Labour Party speech, 1992. The dashed line is the Weibull distribution. Reproduced from Tsizhmovska and Martyushev (2021) under the CC-BY license (https://creativecommons.org/licenses/by/4.0/).

The authors’ reasoning behind this particular type of distribution is couched in terms of the Least Effort principle originally proposed by Zipf (1949) as follows:

“…The author strives to express each of his thoughts in the most economical, shortest way. As a result, the author consciously and unconsciously tends to use sentences of the minimum length (L), among the variety of those that are similar in content {L_1_, L_2_, …, L_n_}, i.e., L = min{L_1_, L_2_, …, L_n_}. It is well known from mathematical statistics […] that the distribution of the minima of a random variable corresponds to the Weibull distribution (strictly, if L = min{L_1_, L_2_, …, L_n_}, n → ∞ and, L_1_, L_2_, …, L_n_ being identically distributed random variables equal to zero or larger, L will obey the Weibull distribution function […]). Thus, the principle of least effort unambiguously indicates that sentence lengths, with a sufficiently large sample, should be described by the Weibull distribution. [p.2]

Other studies in corpus linguistics demonstrate that the dependency holds for word length in terms of number of letters as well as sentence lengths in terms of number of words and can be modeled by a version of the generalized gamma distribution, in particular, gamma [11][12], Weibull [13], or log-normal [14] distributions. All three observed distributions are characterized by similar shape and scale parameters, and choosing among them is not a trivial task [15]. However, all of them capture the exponential-like character of dependency between frequency and length of utterances. In this work we will not take a stand concerning the selection between these particular distributions, and we will use these previous results as an empirical guideline regarding the type of distribution we can expect from the dolphin clicks if they indeed show signs of a communicative system.

### Menzerath-Altmann’s law

This law establishes a reverse correlation between the size of a sequence and the size of its constituent parts. In other words, the longer a sequence, the shorter its constituents, and vice versa. In the realm of natural language, the law applies at different levels of linguistic description, e.g. sentences (size measured in terms of number of clauses), words (measured in terms of number of syllables) and other linguistic units. Altmann (1980) proposed the following formula describing the law:

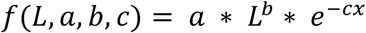

where *f* is the size of a constituent, *L* is the size of the entire sequence, and *a,b,c* are the parameters.

Similarly to Zipf’s laws above, the Menzerath-Altmann law has been shown to hold beyond the domain of natural language, in particular in analysis of genomic sequences [33]. From the information theoretic perspective, both laws are commonly seen as realizations of the tendency for compression or minimization of the linguistic code without the loss of information [16]. Compression was even suggested as a universal principle guiding animal behavior in general [16], as well as communication in particular, although, with regard to the latter, the results were somewhat mixed and dependent on the species and selected unit of analysis [17][18][19][20].

## Method

### Recordings

In a series of six underwater recording sessions that took place between November 2019 and February 2021, we recorded vocalizations freely occurring within the family of five Caribbean bottlenose dolphins (2 males, 2 females and 1 male infant) living in the controlled aquatic environment at the Varna Dolphinarium (Varna, Bulgaria). The characteristics of the recording equipment and recording parameters used in the sessions are described in Supplementary material. Each recording session lasted 5 minutes. During the recording the dolphins were freely swimming, not distracted or engaged in any human-initiated activity. Although our recording setup did not allow for the possibility of attributing specific acoustic productions to particular dolphins, it was ensured that collected sounds were not distorted by distances from the signal receiver (hydrophone) or other extraneous factors.

### Signal analysis

The raw recorded 5 minute-long vocalization streams were denoised via digital 4^th^-order IIR Butterworth high-pass filter with cut-off frequency of 100 Hz. The sequences of clicks were extracted from the records using the short-time (frames of 100 ms) median frequency: any signal frame with median frequency above 5 kHz and maximum level above 0.05 was considered as active. Extracted sequences were amplitude normalized. Individual clicks were identified via envelope detection using the following routine: (i) filtration with 2^nd^-order IIR Butterworth high-pass filter with cut-off frequency of 5 kHz; (ii) sequence segmentation with frame duration of 2.5 ms; (iii) coarse envelope detection via short-time kurtosis computation – every frame with kurtosis above 30 is marked as one containing an individual click; (iv) finally, all individual peaks are processed further in order to remove any possible reflection/reverberation (see Figures 4 and 5).

**Figure 4.**
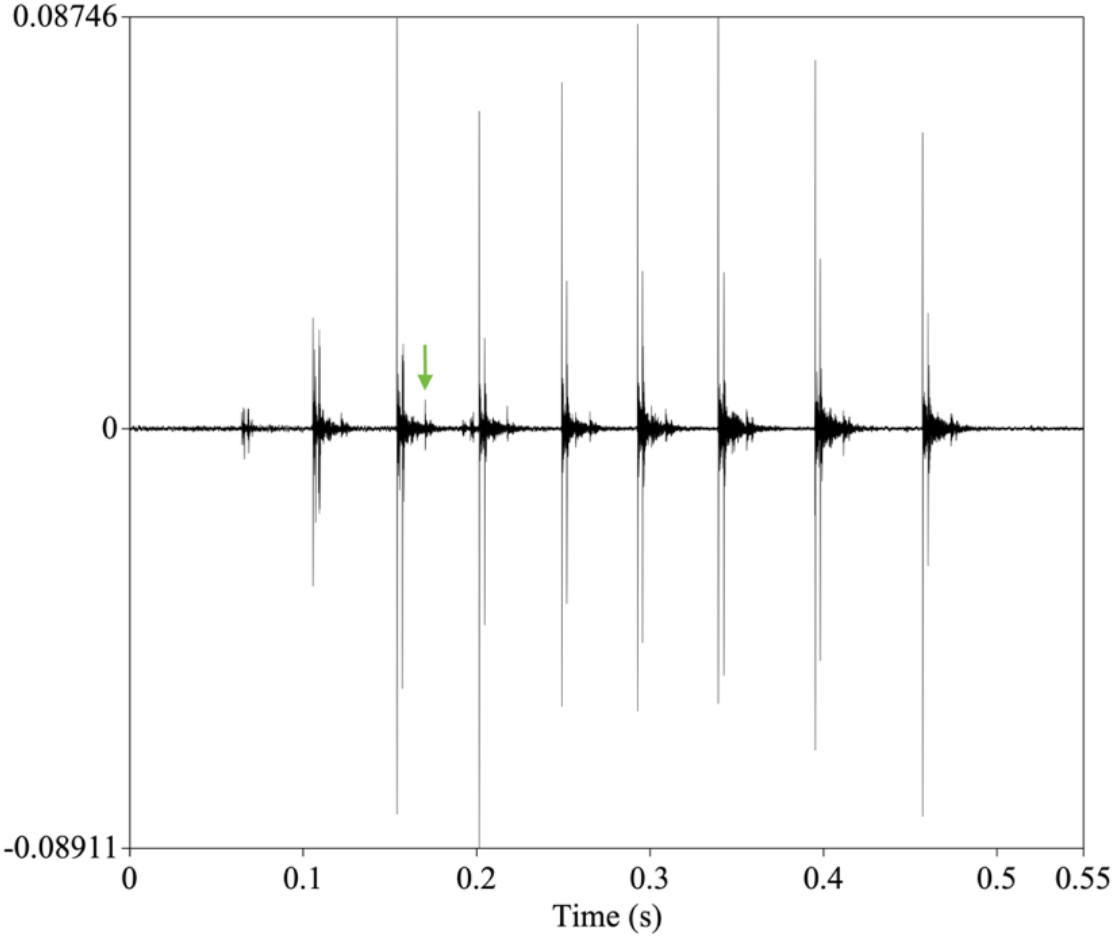
A typical sequence of 9 clicks, selected for the analysis. The pulses in the sequence seem to follow a broadly similar structure. The green arrow points to a possible echo from a reflected pulse.

**Figure 5.**
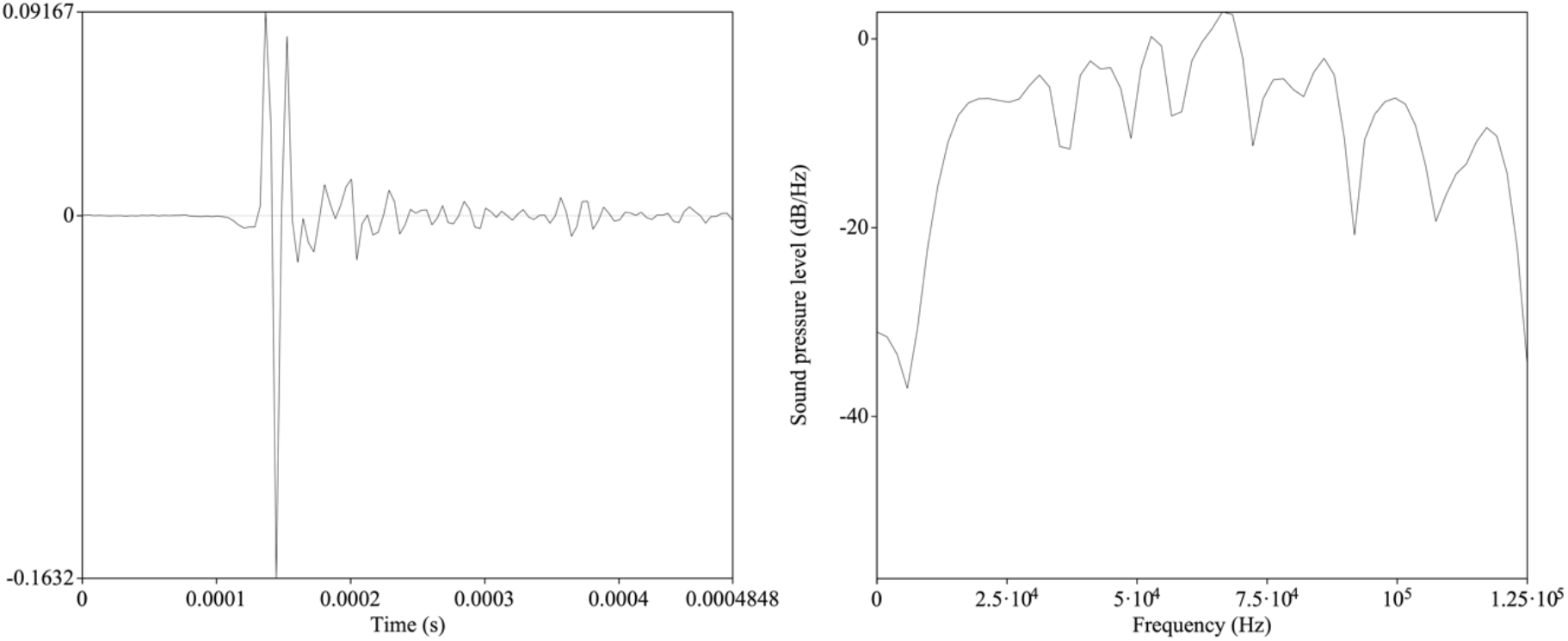
The structure of a typical pulse in the sequence (left) and the distribution of its frequency spectrum (right).

The amplitude contour for each candidate pulse was then sampled into 37 equal temporal bins of the size of 4 µs each. Correspondingly, in the final dataset, each click is represented as a 1 × 37 vector of amplitude values which can be either positive or negative. We also calculated the number of clicks constituting each sequence (sequence size), overall sequence durations as well as durations of individual clicks estimated as an interval between two click onsets within a single click sequence.

### Statistical analysis

We categorized the collected dataset of clicks into natural classes or types using the affinity propagation (AP) method of cluster analysis [21]. This method has an advantage over a more traditionally used k-means cluster analysis in that there is no need to postulate the number of clusters *ad hoc*. Rather, by viewing each data point as a node in a network, the AP algorithm takes measures of similarity between pairs of data points as input and recursively transmits real-valued ‘messages’ along edges of the network until a good set of clusters emerges alongside with final exemplars. The algorithm allows one to control sensitivity or granularity of the clustering process via specifying the ‘input preference’ parameter, or a particular quantile of the input similarities. Using the ‘apcluster’ package in R [22], we first computed a similarity matrix for the amplitude data in the final dataset (see above). Negative squared errors (Euclidean distance) were used as the similarity measure. Erring on the conservative side, we set the initial input preference parameter value at 0.8 which calls for a higher degree of similarity and potentially higher number of clusters. We take the cluster size to be a good approximation for the occurrence frequency of the corresponding click type.

To investigate whether the length of the sequence has an effect on the frequency of use along the lines of Zipf’s law of brevity, we fitted the observed sequence length distribution against theoretical gamma, Weibull and log-normal distributions using the *fitdistrplus* package in R. The fits were evaluated via the Kolmogorov–Smirnov (K-S) criterion whereby a larger p-level value indicates a better fit, as well as by a visual inspection of the respective cumulative distribution function plots.

To analyze a possible relation between the length of a click sequence and duration of its components along the lines predicted by the Menzerath-Altmann law we fitted linear mixed effect models using the *lme4* package [23] in R v. 4.0.2 [24]. Click duration was included as a response variable and sequence length as the only fixed factor. In addition, sequence code and recording session were included as random effects to account for repeated measurements. The original distribution of click durations showed a slight positive skew. Consequently, the dependent variable was square-root transformed for the purposes of the model; the model residuals were visually inspected for homoscedasticity and normal distribution via the *qqplot* and their distribution against the fitted values. We used the likelihood-ratio test to evaluate the significance of the full model which included the fixed and random factors against the null model that included the random factors only. 95% confidence intervals (CIs) and p-values for obtained estimates were computed using the Wald approximation.

## Results

Overall, we identified 3618 clicks organized in 98 sequences. The clicks were produced in the broadband frequency range of 25÷125 kHz which is consistent with previous reports [1][2][3]. The length of sequences varied between 6 and 257 clicks (*median*=23, *IQR*=14-48) and their duration varied within the range of 178-9914 ms (*median*=992, *IQR*=615-1656). Duration of individual clicks varied in the range of 0.37-490 ms (*median*=36, *IQR=*27-47).

The clustering algorithm applied to the array of amplitude vector representations of individual clicks converged after 543 iterations yielding a total of 848 clusters or click categories. The size of the resulting clusters varied between 1 and 120 click tokens. A linear regression mapping the log rank of the signal against log frequency of occurrence (=log size of the cluster) showed a slope of −0.94 (Figure 6) indicating that the distribution of clicks in our tested dolphins’ repertoire is non-random, and furthermore, approximating that in human language samples (see above) in accord with Zipf’s rank-frequency law.

**Figure 6.**
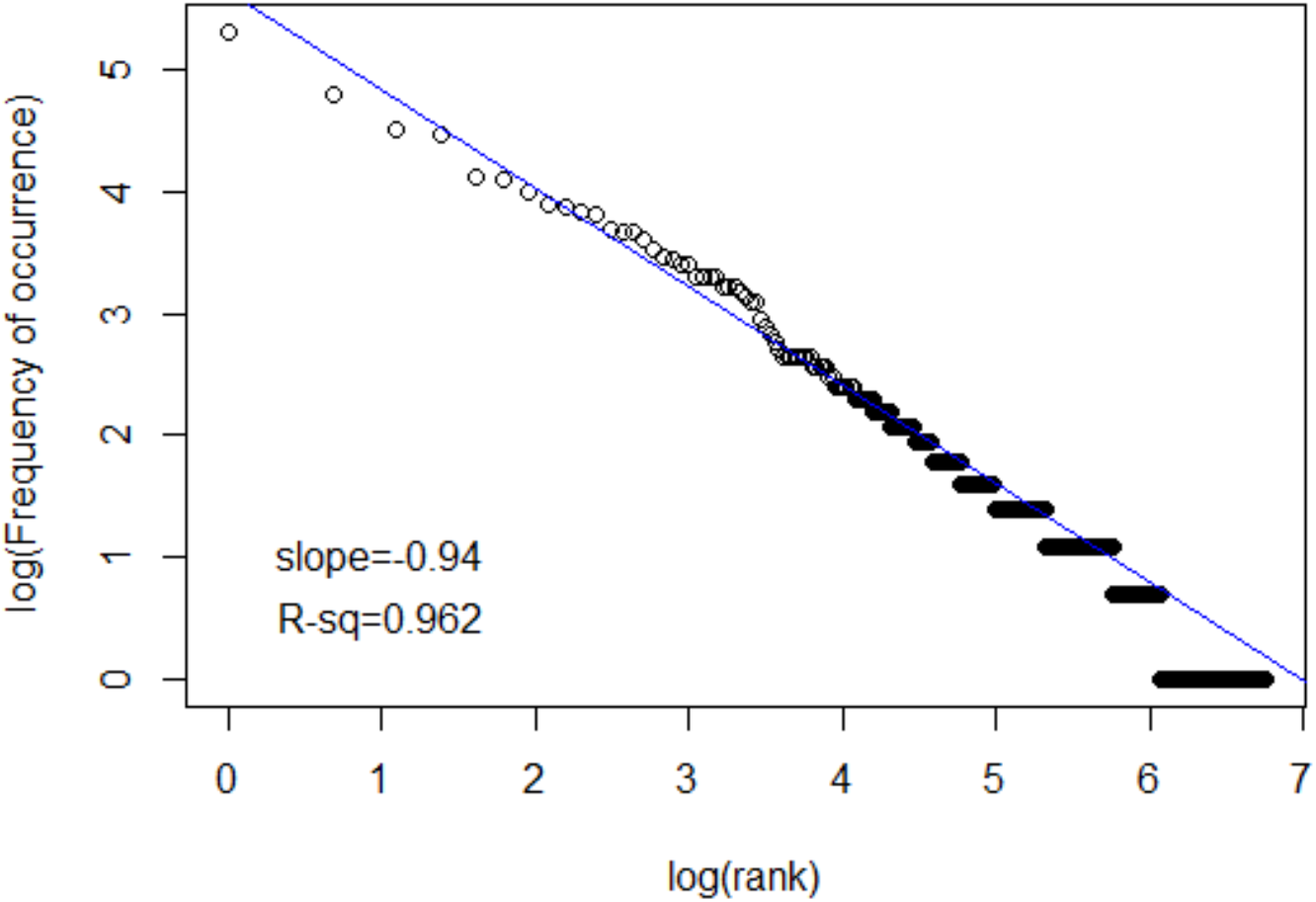
Regression of log (rank) versus log (frequency of occurrence) and lines of best fit for the click repertoires of adult dolphins

Figure 7 shows a probability density plot of sequence lengths in terms of number of pulses across the entire dataset and the fit curves, along with the corresponding cumulative distribution (CDF) functions. We found that the distribution of sequence lengths indeed receives a good fit by the family of the generalized gamma distributions, in particular, a gamma (K-S goodness-of-fit test: *D*=0.118, *p*=0.1321), Weibull (K-S: *D*=0.10248, *p*=0.2548) and log-normal (K-S: *D*=0.072666, *p*=0.6788) distributions. The K-S criterion and the visual inspection of the CDF functions suggests that our click data are best fit by the log-normal distribution.

**Figure 7.**
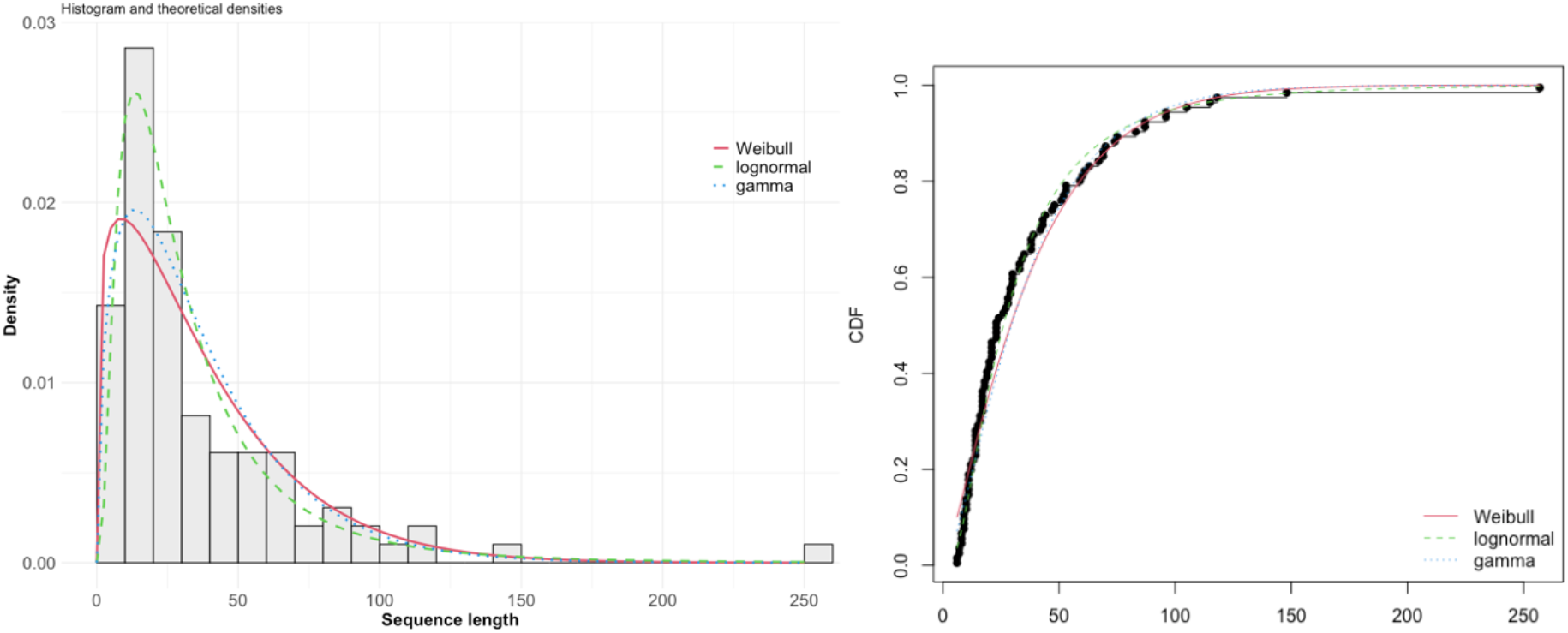
Distribution density of sequence lengths calculated by number of pulses in the analyzed corpus of (left) and the respective cumulative distribution function (right), along with the fit curves.

Finally, we found that sequence length had a statistically significant (model full vs. null: 𝒳^2^(1)=65.79, *p* < 0.001) and negative effect on the click duration (beta= −1.03e-04, 95% CI[−1.28e-04, −7.85e-05], *t*(3448)=-8.18, *p*<.001). In terms of the untransformed dependent variable, increasing a sequence length by one click leads to decrease of duration of individual clicks by about 0.4 ms.

## Discussion

Our results indicate that the repertoire of broadband click sounds produced by our tested dolphins conforms to three key linguistic laws. The number of occurrences of particular (type of) clicks in the dataset as identified by the clustering algorithm inversely correlated with their frequency rank. This is consistent with the idea that, similarly to human languages, this repertoire is formed as a result of two opposing selectional pressures, one aiming for a greater unification and reduced number of repetitions and the other aiming for a greater token diversity. The observed slope value of −0.94 is on a par with that previously reported in the domain of dolphins’ whistles [8][9][10]. This might indicate a similar strategy of use of both signal types in communication, which is conceivable given their common anatomical source, namely, nasal or phonic lips [1][2][3][30][31]. Distribution of sequence lengths followed a log-normal distribution typical for natural language in accord with the law of brevity. Furthermore, the length of a click sequence was shown to inversely correlate with the duration of particular clicks, in accord with Menzerath-Altmann’s law.

If clicks are used for communicative purposes in ways predicted by Zipf’s law series, one can expect some level of structural sophistication in the manner click sequences are formed to express meanings in the dolphins’ putative language. We did not control the contextual variables in our study so it was not possible to correlate specific clicks or click sequences with possible meanings they could be conveying (cf. [25]). Nevertheless, the structural complexity of utterances can be fruitfully studied independently of associated meanings [26]. One pertinent question is what levels of linguistic building blocks are represented by individual clicks and click sequences. Human language is characterized by the ‘duality of patterning’ whereby meaningless units such as phonemes are combined together to produce minimal meaning-bearing conglomerates, e.g. morphemes, and further up the structural hierarchy reaching ultimately the level of a complete message or utterance [27]. In the realm of animal communication, duality of patterning has not so far been attested in its entirety, but some aspects of it were attested in certain species [28]. The sheer variety of click types identified in our study suggests that individual clicks may be some analogues of words, or units conventionalized for use with a fixated lexical or functional meaning. Under this hypothesis, of particular interest are potential syntactic rules that are used in concatenating the clicks into sequences [28][29]. Investigation of these syntactic rules requires a fine-grained analysis of conditional probabilities of occurrence of various click combinations, possibly along the lines of entropy computation that has been suggested for dolphins’ whistles [10]. We also stress that the click types in our study were categorized statistically rather than experimentally. Categorization, e.g. categorical perception, experiments should be conducted to confirm or modify our hypotheses concerning the established click categories.

## Acknowledgements and funding

The authors thank the company ‘Festa Dolphinarium Varna Ltd.’ This work was supported by the European Regional Development Fund within the OP ‘Science and Education for Smart Growth 2014-2020’, project No. BG05M2OP001-1.002-0023.

## Data Acquisition and Signal Detection Parameters

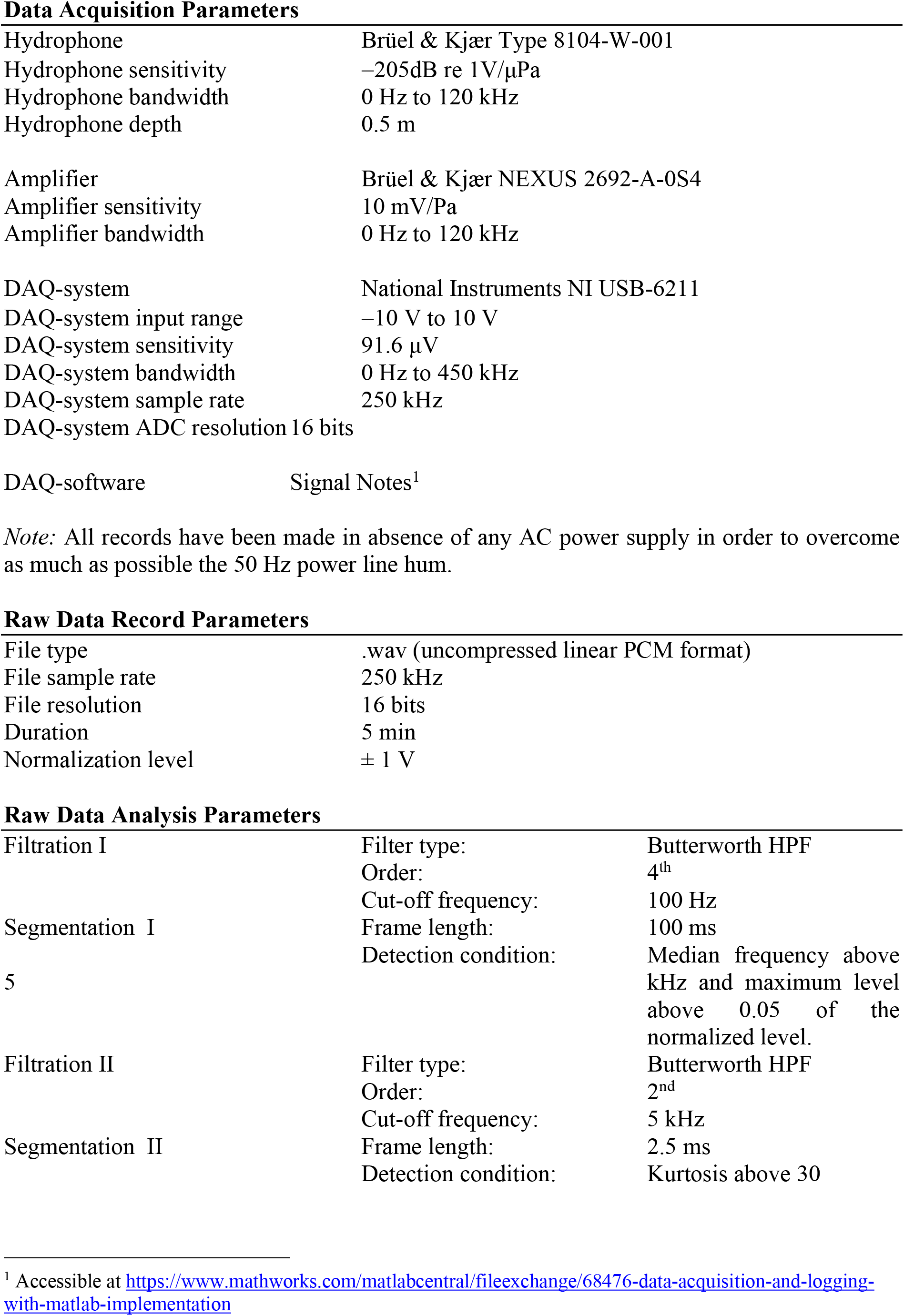

|  |  |
| --- | --- |
| Hydrophone | Brüel & Kjær Type 8104-W-001 |
| Hydrophone sensitivity | −205dB re 1V/μPa |
| Hydrophone bandwidth | 0 Hz to 120 kHz |
| Hydrophone depth | 0.5 m |
| Amplifier | Brüel & Kjær NEXUS 2692-A-0S4 |
| Amplifier sensitivity | 10 mV/Pa |
| Amplifier bandwidth | 0 Hz to 120 kHz |
| DAQ-system | National Instruments NI USB-6211 |
| DAQ-system input range | −10 V to 10 V |
| DAQ-system sensitivity | 91.6 μV |
| DAQ-system bandwidth | 0 Hz to 450 kHz |
| DAQ-system sample rate | 250 kHz |
| DAQ-system ADC resolution | 16 bits |
| DAQ-software | Signal Notes<sup>1</sup> |

*Note:* All records have been made in absence of any AC power supply in order to overcome as much as possible the 50 Hz power line hum.

**Raw Data Record Parameters**
|  |  |
| --- | --- |
| File type | .wav (uncompressed linear PCM format) |
| File sample rate | 250 kHz |
| File resolution | 16 bits |
| Duration | 5 min |
| Normalization level | ± 1 V |

**Raw Data Analysis Parameters**
|  |  |  |
| --- | --- | --- |
| Filtration I | Filter type: | Butterworth HPF |
|  | Order: | 4<sup>th</sup> |
|  | Cut-off frequency: | 100 Hz |
| Segmentation I | Frame length: | 100 ms |
| 5 | Detection condition: | Median frequency above kHz and maximum level above 0.05 of the normalized level. |
| Filtration II | Filter type: | Butterworth HPF |
|  | Order: | 2<sup>nd</sup> |
|  | Cut-off frequency: | 5 kHz |
| Segmentation II | Frame length: | 2.5 ms |
|  | Detection condition: | Kurtosis above 30 |

<sup>1</sup> Accessible at https://www.mathworks.com/matlabcentral/fileexchange/68476-data-acquisition-and-logging-with-matlab-implementation

